# Predicting the presence and titer of rabies virus neutralizing antibodies from low-volume serum samples in low-containment facilities

**DOI:** 10.1101/2020.04.24.060095

**Authors:** Diana K. Meza, Alice Broos, Daniel J. Becker, Abdelkader Behdenna, Brian J. Willett, Mafalda Viana, Daniel G. Streicker

## Abstract

Serology is a core component of the surveillance and management of viral zoonoses. Virus neutralization tests are a gold standard serological diagnostic, but requirements for large volumes of serum and high biosafety containment can limit widespread use. Here, focusing on *Rabies lyssavirus,* a globally important zoonosis, we developed a pseudotype micro-neutralization rapid fluorescent focus inhibition test (pmRFFIT) that overcomes these limitations. Specifically, we adapted an existing micro-neutralization test to use a green fluorescent protein–tagged murine leukemia virus pseudotype in lieu of pathogenic rabies virus, reducing the need for specialized reagents for antigen detection and enabling use in low-containment laboratories. We further used statistical analysis to generate rapid, quantitative predictions of the probability and titer of rabies virus neutralizing antibodies from microscopic imaging of neutralization outcomes. Using 47 serum samples from domestic dogs with neutralizing antibody titers estimated using the fluorescent antibody virus neutralization test (FAVN), pmRFFIT showed moderate sensitivity (78.79%) and high specificity (84.62%). Despite small conflicts, titer predictions were correlated across tests repeated on different dates both for dog samples (r = 0.93), and for a second dataset of sera from wild common vampire bats (r = 0.72, N = 41), indicating repeatability. Our test uses a starting volume of 3.5 μL of serum, estimates titers from a single dilution of serum rather than requiring multiple dilutions and end point titration, and may be adapted to target neutralizing antibodies against alternative lyssavirus species. The pmRFFIT enables high-throughput detection of rabies virus neutralizing antibodies in low-biocontainment settings and is suited to studies in wild or captive animals where large serum volumes cannot be obtained.

## Introduction

The last few decades have seen a surge in newly emerging human viruses that originate from wildlife (Cunningham, Daszak, & Wood, 2017; Daszak, Cunningham, & Hyatt, 2000; Goodin, Jonsson, Allen, & Owen, 2018). Key examples include Nipah virus (Gurley et al., 2017), Seoul virus (Kerins et al., 2018), 2009 H1N1 (Mena et al., 2016), and the recent SARS-CoV-2 (Zhou et al., 2020). Understanding the epidemiological dynamics of such viruses within their natural host populations is a fundamental component to discerning the spatiotemporal dynamics of past outbreaks and anticipating future emergence (Cunningham et al., 2017; Plowright et al., 2016). Investigating the dynamics of zoonotic viruses within wildlife presents multiple challenges such as limited sample sizes, biased sampling and multiple diagnostic tests that are difficult to compare. Moreover, viruses themselves may be undetectable at the moment of sampling when infections periods are short or virus shedding is intermittent (Becker, Crowley, Washburne, & Plowright, 2019; Gilbert et al., 2013; Plowright, Becker, McCallum, & Manlove, 2019). Consequently, key parameters needed to inform population-level disease dynamics (e.g. incidence, infection and incubation periods) are difficult or impossible to measure directly (Borremans, Hens, Beutels, Leirs, & Reijniers, 2016). Serological tests offer a powerful alternative to approaches that rely on pathogen detection (Gilbert et al., 2013). Since pathogen-specific antibodies generally persist longer than the pathogen itself, serological data can inform individual exposure histories, population seroprevalence, and in some cases, help approximate the force of infection (Borremans et al., 2016; Gamble, Garnier, Chambert, Gimenez, & Boulinier, 2020; Gilbert et al., 2013; Metcalf et al., 2016). However, applying serology to longitudinal studies of wildlife presents distinct challenges from tests used in clinical diagnostic settings, where accuracy takes precedence over scalability. In particular, serological tests for studies of wildlife should be amenable to the small volumes of serum that often characterize collections from small-bodied hosts (e.g. rodents, bats, birds), scalable to large numbers of samples required for such studies, and possible to implement in low-biocontainment laboratories.

*Rabies lyssavirus* (RV; Genus *Lyssavirus,* Family *Rhabdoviridae)* is a zoonotic virus transmitted in saliva by the bite of infected mammals (mainly Carnivora and Chiroptera) (Rupprecht, Kuzmin, & Meslin, 2017). RV constitutes a substantial global health problem that is responsible for over 59,000 human fatalities annually, mostly attributable to domestic dogs (WHO, 2017). In Latin America, vampire bat RV causes more cases of human and domestic animal rabies mortality than RV transmitted by dogs; similarly, bats are among the most frequently diagnosed sources of human rabies exposure in the USA (Ma et al., 2020; Schneider et al., 2009). Bat lyssaviruses exemplify the challenges of studying zoonoses using pathogen detection and the potential advantages of serological inference (Turmelle et al., 2010). Active infections are rarely detected due to low population-level incidence and the short infectious period that precedes death (Jackson et al., 2008; Turmelle et al., 2010). However, abortive infections (i.e. bats that are exposed to RV but do not become infectious) are routinely observed through antibody presence in apparently healthy bats (Constantine, Tierkel, Kleckner, & Hawkins, 1968; Jackson et al., 2008; Obregón-Morales et al., 2017; Steece & Altenbach, 1989). As such, antibody detection is commonly used for ecological and epidemiological studies of bat rabies (Blackwood, Streicker, Altizer, & Rohani, 2013; Costa et al., 2013; de Thoisy et al., 2016; George et al., 2011; Horton et al., 2020; Streicker et al., 2012; Wright et al., 2010).

Existing tests to detect RV antibodies include enzyme-linked immunosorbent assays (ELISAs) (Barton & Campbell, 1988; Cliquet et al., 2004; Wasniewski & Cliquet, 2012) and virus neutralization tests (Cliquet, Aubert, & Sagné, 1998; Smith, Yager, & Baer, 1973). ELISAs are a rapid and effective method for testing large numbers of samples, but they do not detect virus neutralization (Ma, Niezgoda, Blanton, Recuenco, & Rupprecht, 2012; Welch, Anderson, & Litwin, 2009). Moreover, since ELISAs ideally require a secondary antibody specific to the host species of the original sample (Reynes et al., 2004), which is often non-existent, they are not widely used in wildlife rabies surveillance. Tests detecting virus neutralizing antibodies (VNA) are more commonly adopted (De Benedictis, Mancin, Cattoli, Capua, & Terrregino, 2012; Irie & Kawai, 2018; Moore & Hanlon, 2010). Specifically, the fluorescent antibody virus neutralization test (FAVN) and the rapid fluorescent focus inhibition test (RFFIT) are considered the gold standard for measuring vaccination response by the World Health Organization (WHO) (Briggs et al., 1998; De Benedictis et al., 2012). Both the FAVN and RFFIT have been modified to facilitate use with wildlife samples. For instance, Kuzmin et al. (2008) modified the RFFIT into a micro-neutralization test using 4-well Teflon coated slides instead of 8-well chamber slides. This produced a sensitive and specific test that only required 3.5 μl of serum (regular RFFIT and FAVN require ~50 μL of serum) (Kuzmin et al., 2008). Several other laboratories have introduced the modification of pseudotype viruses (i.e. non-rabies viruses engineered to express heterologous envelope glycoproteins) to quantify VNA titers without highly pathogenic live lyssaviruses (Temperton, Wright, & Scott, 2015), which in most countries involves biosafety level (BSL)-3 laboratories to produce concentrated virus stocks (WHO, 2018). Pseudotyped viruses based on vesicular stomatitis virus (Moeschler, Locher, Conzelmann, Krämer, & Zimmer, 2016), human immunodeficiency virus, and murine leukemia virus (MLV) (Wright et al., 2008) tend to be highly sensitive and specific relative to live lyssavirus counterparts. Viral pseudotypes have also been modified to incorporate molecular biomarkers such as firefly luciferase, renilla luciferase, or green fluorescent protein (GFP), eliminating the need to fix and stain cells with a fluorescein isothiocyanate (FITC) conjugated rabies antibody, which reduces test cost (Bentley, Mather, & Temperton, 2015; Mather et al., 2013; Moeschler et al., 2016; Moore & Hanlon, 2010; Wright et al., 2008).

Despite these improvements, existing variations of RV neutralization tests have several limitations. First, all tests require serial dilutions of serum samples, each of which requires manual microscope scoring that increases personnel time and risks of error (Moeschler et al., 2016; Péharpré et al., 1999). Second, testing is commonly split into screening and end-point titration phases, requiring tests to be run in duplicate or sequentially. Third, pseudotypes have not yet been integrated into microneutralization tests, implying that tests to detect VNAs against most lyssaviruses in their natural bat reservoirs can only be carried out in BSL-3 or higher facilities. The growing demand to study wildlife on large spatial and temporal scales would benefit from minimizing these logistical constraints (Becker et al., 2019).

The ideal serological diagnostic would be a scalable virus neutralization test that requires low-volume samples and can be carried out in any standard microbiology/cell culture laboratory. Here, the microneutralization RFFIT is adapted to use an MLV-based viral pseudotype bearing the RV glycoprotein and carrying a GFP marker gene. Further, we introduce a novel quantitative approach that combines digital image analysis of infected cells with statistical analysis, which allows us to estimate rabies virus neutralizing antibody (RVNA) titers without multiple serum dilutions or rounds of testing. Our approach relies on the expectation of a negative relationship between RVNA concentrations in serum and the number of cells that become infected upon viral challenge in the presence of that serum. Defining that relationship quantitatively using known titers of standard rabies immune globulin (SRIG) allows predicting the presence and, when appropriate, titers of RVNAs in test sera. Our approach, which we refer to as a pseudotype micro-neutralization RFFIT (hereafter, pmRFFIT, **Figure 1**) provides a safe, low-cost alternative to standard RV neutralization tests that is suitable for large-scale, population-level studies or laboratory studies where small animals are longitudinally sampled.

**Figure 1.**
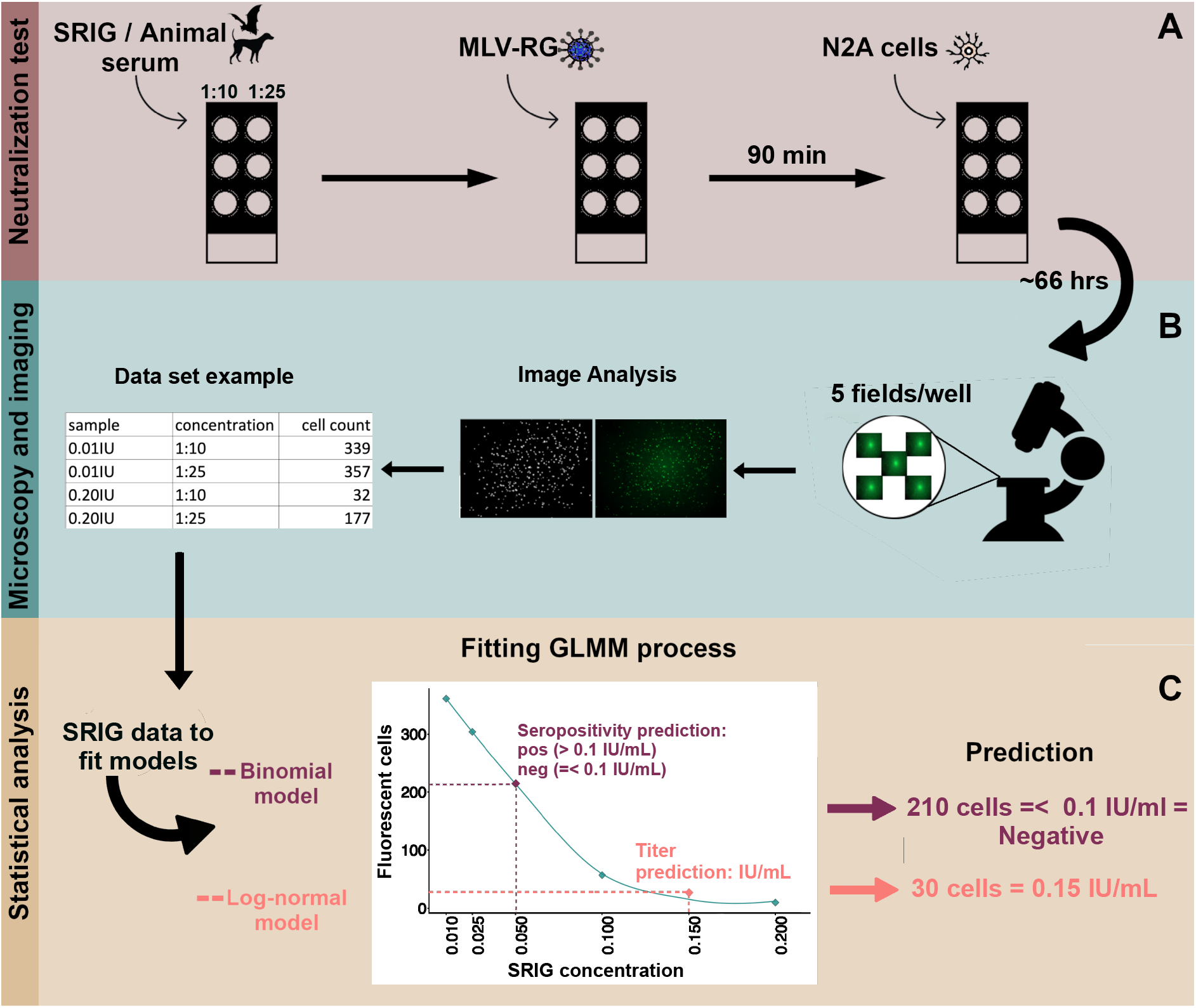
Workflow of the pmRFFIT approach. A) Set up of the neutralization test using an MLV(RG) pseudotype and 2 dilutions of either SRIG or animal serum. B) Microscopy phase and imaging to perform the cell count of the fluorescent cells to construct a database to fit the statistical models. C) Construction of the statistical models with two different types of prediction.

## Methods

### Laboratory work

#### Cell lines

For all the neutralization tests, mouse neuroblastoma cells, N2A (Neuro-2A, ATCC^®^ CCL131™), were cultured in Minimum Essential Media (MEM) supplemented with 10% fetal bovine serum (FBS), 1% 100x non-essential amino acid (NEAA), 1% 200mM L-glutamine and 1% antibiotic-antimycotic. For viral pseudotype production, human embryonic kidney 293T cells (293T, ATCC^®^ CRL-3216™), were cultured in Dulbecco’s Modified Eagle’s Medium (DMEM) supplemented with 10% FBS, 1% 200mM L-glutamine, 100 units/mL Penicillin, 1 μg/ml Streptomycin and 400 μg/mL G418 (Geneticin) antibiotic.

#### Pseudotype production

MLV pseudotypes expressing the challenge virus standard-11 (CVS-11) rabies virus glycoprotein (MLV(RG)) were generated by co-transfection of pCMVi (MLV gag-pol expression vector), pCNCG (MLV viral origin and GFP reporter gene), and pI.18-CVS-11 (plasmid encoding the CVS-11 RG) (Bock, Bishop, Towers, & Stoye, 2000; Towers et al., 2000; Wright et al., 2008). Plasmids were transfected using polyethylenimine (Polysciences, Inc.) into 293T cells at a ratio of 1:1.5:1 of pCMVi/pCNCG/pl.18-CVS-11. After 72 hours, the supernatant containing the viral pseudotypes was harvested, aliquoted and stored at −80°C until further use. Virus concentration was determined in N2A cells by calculating the TCID50 (50% tissue culture infective dose) using the end point method and using the Spearman-Kärber formula (Condit, 2013; Hierholzer & Killington, 1996).

#### Neutralization test

A two-dilution micro-neutralization test was established following Kuzmin et al. (2008); however, in place of 4-well Teflon-coated slides, 6-well Teflon-coated slides (Tekdon Inc.) were used, enabling 3 samples to be analyzed per slide. Each serum sample (starting volume of 3.5 μL), was screened at a 1:10 and 1:25 dilution (a total of 2 wells per sample). Serum dilutions were inoculated with 12.5 μl of MLV(RG) (viral input: 300-400 TCID50) and incubated in humidified square petri dishes at 37°C, 5% CO_2_ for 90 min. After incubation, 25 μl of N2A cells (2×10^6^ cells/ml) were added to the serum-virus mixture and slides were incubated at 37°C, 5% CO_2_ for 66 to 72 h in the humidified square petri dishes. This extended incubation period (compared to the FAVN (48 h) or the RFFIT (20 to 24 h)) was required for the MLV(RG) to generate sufficient GFP expression for later microscopy. Since our pseudotype virus used GFP to indicate viral entry into cells, neither acetone fixation nor staining with anti-rabies monoclonal globulin were required. Every neutralization test included three internal controls: 1) a cell-only control where no virus or serum was added to the well; 2) a virus control, comprising back titrations of the MLV(RG) with: a) the original concentration of the virus (300-400 TCID50), b) a 1:10 dilution of the virus, and c) a 1:100 dilution of the virus; and 3) a negative control for neutralization, where we added 300-400 TCID50 of MLV(RG) to two wells without any serum. A standard curve was generated with each neutralization test describing the expected number of infected cells versus titer concentration. For this, the same protocol was used with six different titer concentrations of SRIG in lieu of serum samples: 0.01, 0.025, 0.05, 0.1, 0.15 and 0.2 IU/mL (reconstituted at 30 IU/mL in 1.0 mL of nuclease-free water, 2^nd^ WHO International Standard, Human; NIBSC, UK) (**Figure 1** A).

### Imaging and data processing

After incubation, slides were photographed at 4x magnification under a fluorescence microscope (EVOS FL Cell Imaging System). Wells were divided into five equally sized fields (four corners and center) and photographed clockwise from the top left corner, such that the fifth photograph was always the center field. Each photograph was processed through the program ImageJ (version 1.52k (Rasband, 2018)) using the Autolocal Threshold Phansalkar plugin (size radius 15) (part of the Fiji distribution (Schindelin et al., 2012)), where each image was transformed into a binary image (every pixel stored as a single bit). This indicated the presence (white) and absence (black in the background) of GFP fluorescence. Next, the command “Analyze Particles” was used to count the total number of fluorescent cells per field (i.e. infected cells). This command grouped and counted the white neighboring pixels with a predetermined size area and circularity to be a single cell (size area: 5-50 circularity: 0.80-1.0), so counts corresponded to the number of infected cells. Cell count outputs were converted into a standardized spreadsheet using a Python version 3.7.2 script (Python Core Team, 2019) (script available in supplementary materials). At the end of the image processing step, each serum sample was described by 10 data points consisting of the number of the fluorescent cells in each of 5 fields (photographs) in the 1:10 and 1:25 dilutions (**Figure 1** B).

### Statistical analysis

All statistical analyses were executed in *R* (R Core Team, 2018). To predict the probability of RVNA presence, a generalized linear mixed model (GLMM) with a binomial distribution was fit to the binary outcome of the cell counts from the SRIG concentration series, where titers ≤ 0.1 IU/mL were considered RVNA negative and > 0.1 IU/mL were considered RVNA positive. Other serology studies in wildlife have used similar thresholds to detect RVNA (Araujo et al., 2014; Azevedo de Paula Antunes et al., 2017; Campos et al., 2019; Silva et al., 2010). To next predict RVNA titers, a GLMM with a lognormal distribution was fit to the infected cell counts across the SRIG concentration series. For both the binomial and log-normal models, a model was constructed including all data (i.e. from 1:10 and 1:25 dilutions) where two fixed effects were considered: 1) the count of virus-infected N2A cells (scaled to improve model convergence) and 2) the serum dilution level (two factors: 1:10 and 1:25 dilution). Random slope and intercept terms were also considered for the date the test was run (“test date”) to account for observed variation in the relationships between SRIG titers and infected cell counts across dates (**Figure 2**) and for the field number (1 to 5) within each microscope well (“field”) to account for variation in cell counts between fields (the middle field, field 5, had more agglomerated cells in particular). To evaluate whether a simpler, single dilution test produced comparable results, the full dataset was then subset to the 1:10 dilution only. The binomial and log-normal models fit to this data subset included only the fixed effect of the virus-infected N2A cell counts, but the random effects were identical to those explained above (i.e. test date and field). Models were fit using the *‘lme4’* package (Bates et al., 2015). The *‘predict’* function was used to generate the predicted probability that a SRIG concentration or serum sample was seropositive (binomial model) and its corresponding RVNA titer (log-normal model). Predictions per field were averaged to obtain results per sample (**Figure 1** C).

**Figure 2.**
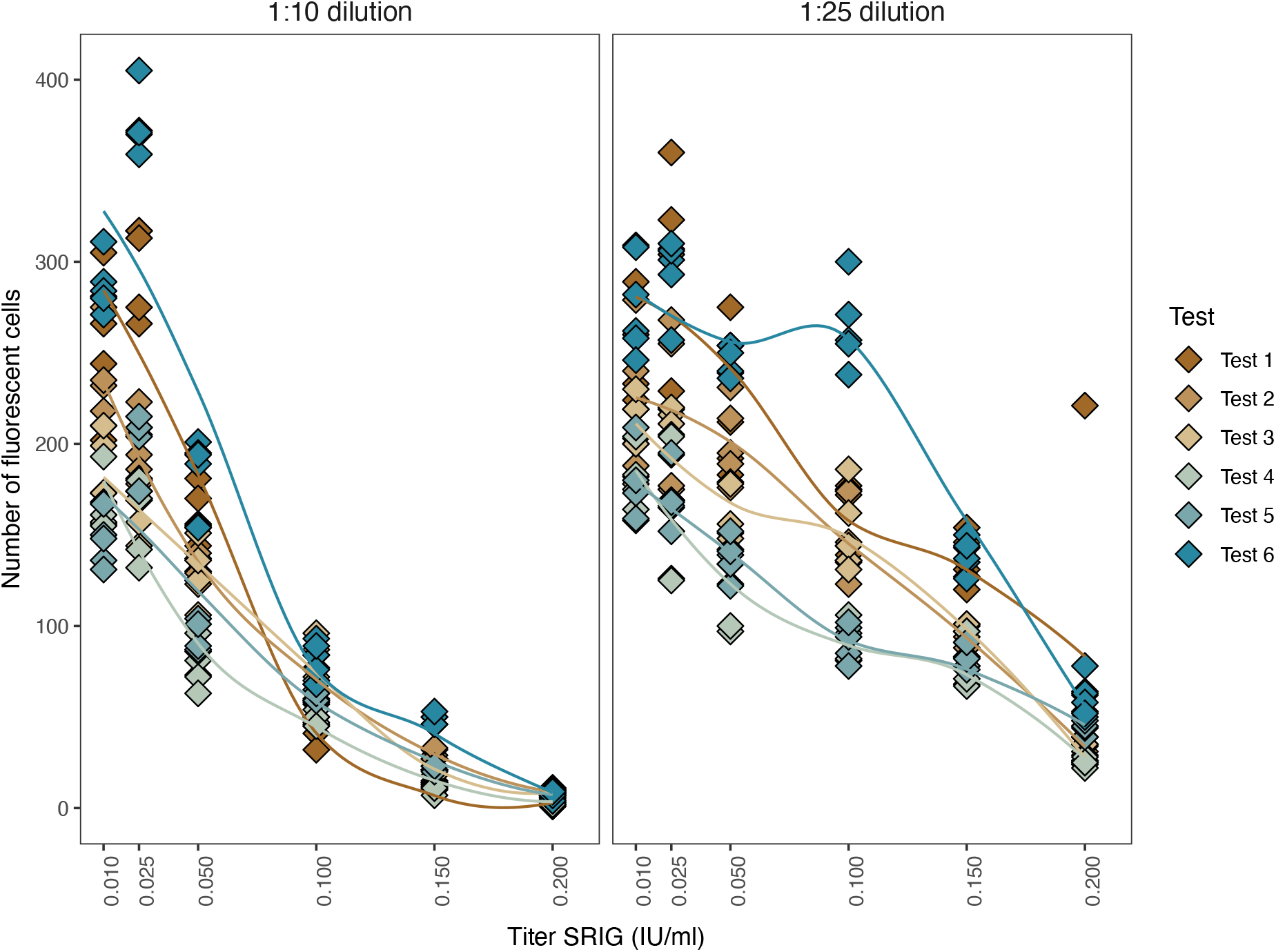
Raw counts of fluorescent cells for the SRIG concentration curves from 6 different pmRFFIT from different test dates and for each dilution. The dilutions were performed on the same date, with one well next to the other. Each color represents a different test date. The loess curves denote differences in the counts from test to test.

#### Model selection

To select the best-performing models, a top-down model selection strategy was implemented (Zuur, Ieno, Walker, Saveliev, & Smith, 2009). First, we identified the optimal random effects by comparing models with alternative random effect structures, but the same fixed effects, using the Akaike Information Criterion corrected for small sample size (AICc) calculated using the *‘MuMIn’* package (Bartoń, 2013). A lower AICc with a difference of at least two values (ΔAICc > 2) was considered as evidence of improved model fit. Next, keeping the chosen random effects, models with the two different datasets were evaluated (simpler model: 1:10 dataset, and more complex model: 1:10 and 1:25 dataset) (**Table 1**). Since models fit to different datasets cannot be compared through AICc, the simpler and more complex models were compared using predictions for SRIG (sera with known titers). For the binomial models, the sensitivity and specificity with a threshold of > 0.1 IU/mL for seropositivity were used. For the log-normal models, the Spearman correlation coefficients of predicted and known SRIG titers were compared.

**Table 1.** Random effect structures investigated for the binomial (B) and log-normal (L) models, and respective model fit measures: AICc, difference in AICc from best model (ΔAICc) and weight (W). The fixed effects for all these models were identical and included the scaled count of the fluorescent cells and the dilution effect

| Model | Random effects | AICc | ΔAICc | W |
| --- | --- | --- | --- | --- |
| **B** | 1. Test date as random intercept and random slope | 91.62 | 0.0 | 0.71 |
|  | 2. Test date as random intercept and random slope + number of field as random intercept | 93.49 | 1.87 | 0.28 |
|  | 3. Test date as random intercept | 102.32 | 10.70 | 0.0 |
|  | 4. Test date as random intercept + number of field as random intercept | 104.28 | 12.66 | 0.0 |
| **L** | 1. Test date as random intercept and random slope + number of field as random intercept | -1746.59 | 0.0 | 0.7 |
|  | 2. Test date as random intercept and random slope | -1744.89 | 1.70 | 0.3 |
|  | 3. Test date as random intercept + number of field as random intercept | -1637.32 | 109.27 | 0.0 |
|  | 4. Test date as random intercept | -1637.30 | 109.29 | 0.0 |

### Test repeatability and external validation

Dog (N = 47) and bat (N = 41) sera were used to assess pmRFFIT repeatability. Dog sera had previously been tested using FAVN at the Animal Health and Veterinary Laboratories Agency, Weybridge (Wright et al., 2009). Bat sera were collected from common vampire bats *(Desmodus rotundus)* as part of an ongoing field study in Peru (collection permit: 0142-2015 SERFOR/DGGSPFFS; exportation permit: 003327-SERFOR; SI 1).

To quantify repeatability, dog and bat sera were processed using the pmRFFIT as described above on two separate dates. For the binomial model, repeatability was measured as the proportion of samples with the same predicted serological status on both test dates. For the log-normal model, repeatability was calculated as the Spearman correlation coefficient of predicted RVNA titers from different test dates.

The sensitivity and specificity of the pmRFFIT were calculated relative to the FAVN using binomial predictions from dog sera. Sera with FAVN titers > 0.1 IU/mL were classified as positive and ≤ 0.1IU/mL as negative. Only data from the first test date were used under the assumption that future applications would preferably use a single test. To compare pmRFFIT accuracy in different FAVN titer ranges, the RVNA values obtained through FAVN were categorized through the *‘cut’* function in R. The number of categories was obtained by dividing the range of the RVNA titer values (maximum minus minimum value obtained through FAVN) by the optimal category width, calculated with the Freedman-Diaconis rule (Freedman & Diaconis, 1981). Later, to evaluate the accuracy of the binomial prediction in the lowest FAVN titers, the categorization procedure was repeated but only using the lowest resulting category from the previous step. A Receiver Operating Characteristic (ROC) curve analysis was used to explore the accuracy of the log-normal predictions across the range of evaluated titers. FAVN titers with a threshold of > 0.1 IU/mL (positives) were used as the benchmark reference.

## Results

To understand the variability of the pmRFFIT, replicate SRIG titer concentration curves were produced on 6 different dates between 30/05/2019 and 27/06/2019 (hereafter “Test 1” through “Test 6”). As expected, the number of infected cells declined at higher SRIG titers in all replicates; however, the shape of the antibody decay curve varied across test dates (**Figure 2**). At the 0.1 IU/mL SRIG concentration, infected cell counts were more dispersed in the 1:25 dilution than in the 1:10 dilution, as indicated by higher interquartile range (IQR) within each of the 6 test dates. Across all the SRIG concentrations in all test dates (N = 36), 77.78% of the count comparisons were less dispersed in the 1:10 dilution suggesting this dilution could be more precise for downstream statistical analysis (SI 2).

### Prediction of seropositivity using binomial GLMM

Binomial GLMMs accurately predicted seropositive and seronegative SRIG concentrations (**Figure 3**). The best random effects for the binomial model included a random slope and intercept for test date (**Table 1**). The models built with the 1:10 dilution data only (“one-dilution model”) and from both the 1:10 and 1:25 dilution data (“two-dilution model”) had equivalent specificity (100%), but the one-dilution model was more sensitive (100% versus 58.33%, **Figure 3** A, B). Furthermore, the two-dilution binomial model failed to correctly predict the seropositive controls on 4 out of the 6 test dates, confirming improved performance of the one-dilution model (**Figure 3** B).

**Figure 3.**
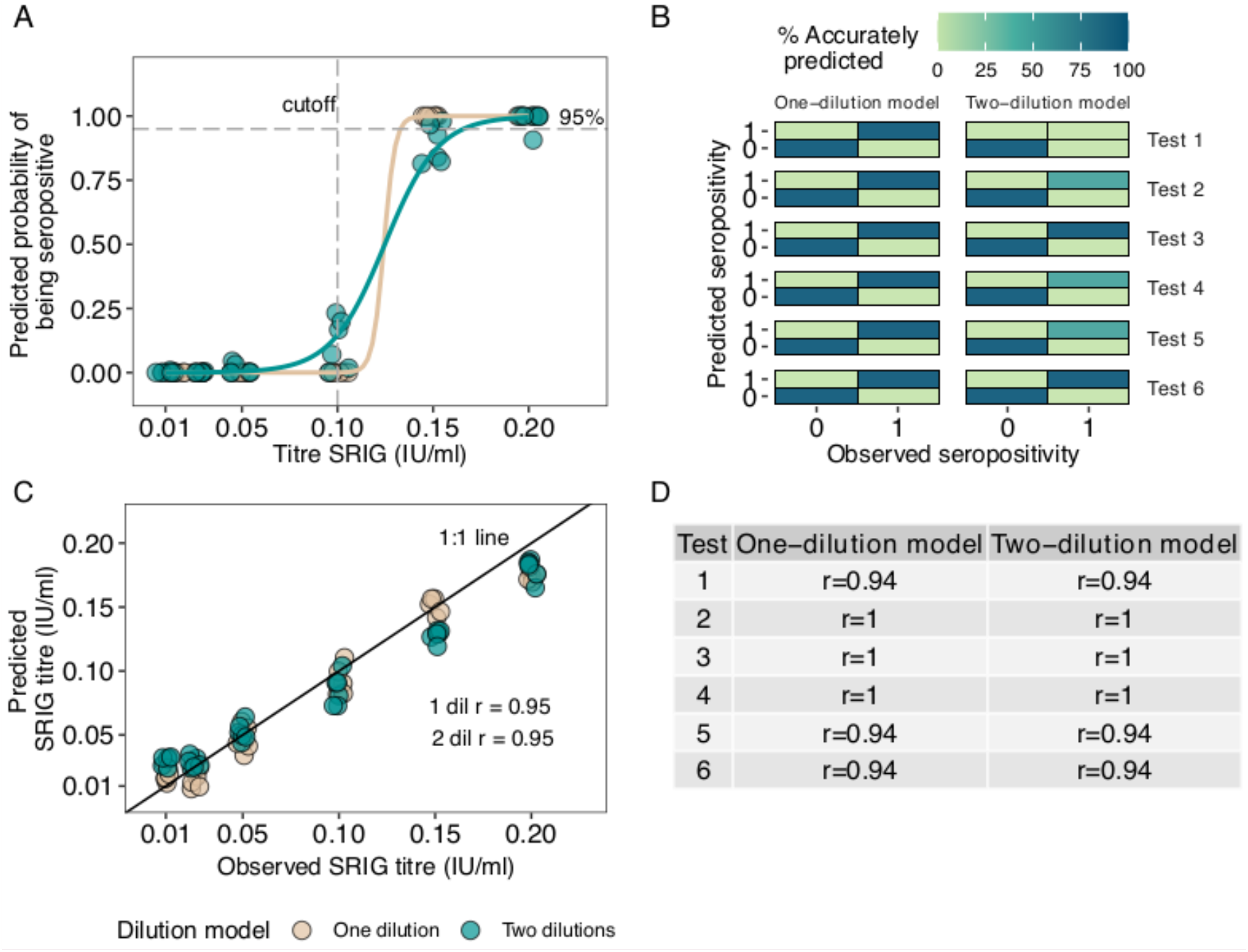
Results of the models fit to the SRIG control data. All graphs compare the one-dilution (1:10) and the two-dilution (1:10 and 1:25) models. A) Model fit of the titers of the SRIG concentration curve versus the predicted probability from the binomial model. The horizontal dash line represents the 95% probability and the vertical dash line represents the previously set threshold at 0.1 IU/mL. B) Sensitivity and specificity of the binomial prediction per test date. C) Correlation of the titers of the observed SRIG values versus the predicted values with the log-normal models. D) Comparison of the Spearman coefficients between the correlations of the observed titers versus the predicted titers per test date.

### Prediction of titer values using log-normal GLMM

The log-normal GLMMs gave repeatable predictions of RVNA titers from the datasets generated through our protocol across test dates (**Figure 3** C). The best log-normal model included a random intercept and slope for test date. Although the most complex model had the lowest AICc, the simpler model (without the random intercept of field) had a ΔAICc < 2 (**Table 1**). Observed and predicted SRIG titers were highly correlated for both the one and two-dilution models (r = 0.95, **Figure 3** C). When comparing test dates (i.e. one-to-one comparison between correlations of the one-dilution and the two-dilution model from the same test dates), the correlation coefficients were similar, suggesting the simpler one-dilution model is sufficient for titer prediction (**Figure 3** D).

### Repeatability and external validation with animal sera

Among all the animal serum samples (N = 88), 95.45% were consistently predicted as seropositive or seronegative (92.68% for bats and 97.87% for dogs). Bat samples (N = 41) had a repeatability of 66.7% for positive samples (6/9 positive samples were positive in the second round) and 100% for negative samples (N = 32). Dog samples (N = 47) had 100% repeatability for positive samples (N = 28) and 94.74% for negative samples (18/19 negative samples remained negative in the second round) (**Figure 4** A, SI 3). Predicted titers between test repetitions were correlated for both sera (both r = 0.81, bat r = 0.72, dog r = 0.93, **Figure 4** B).

**Figure 4.**
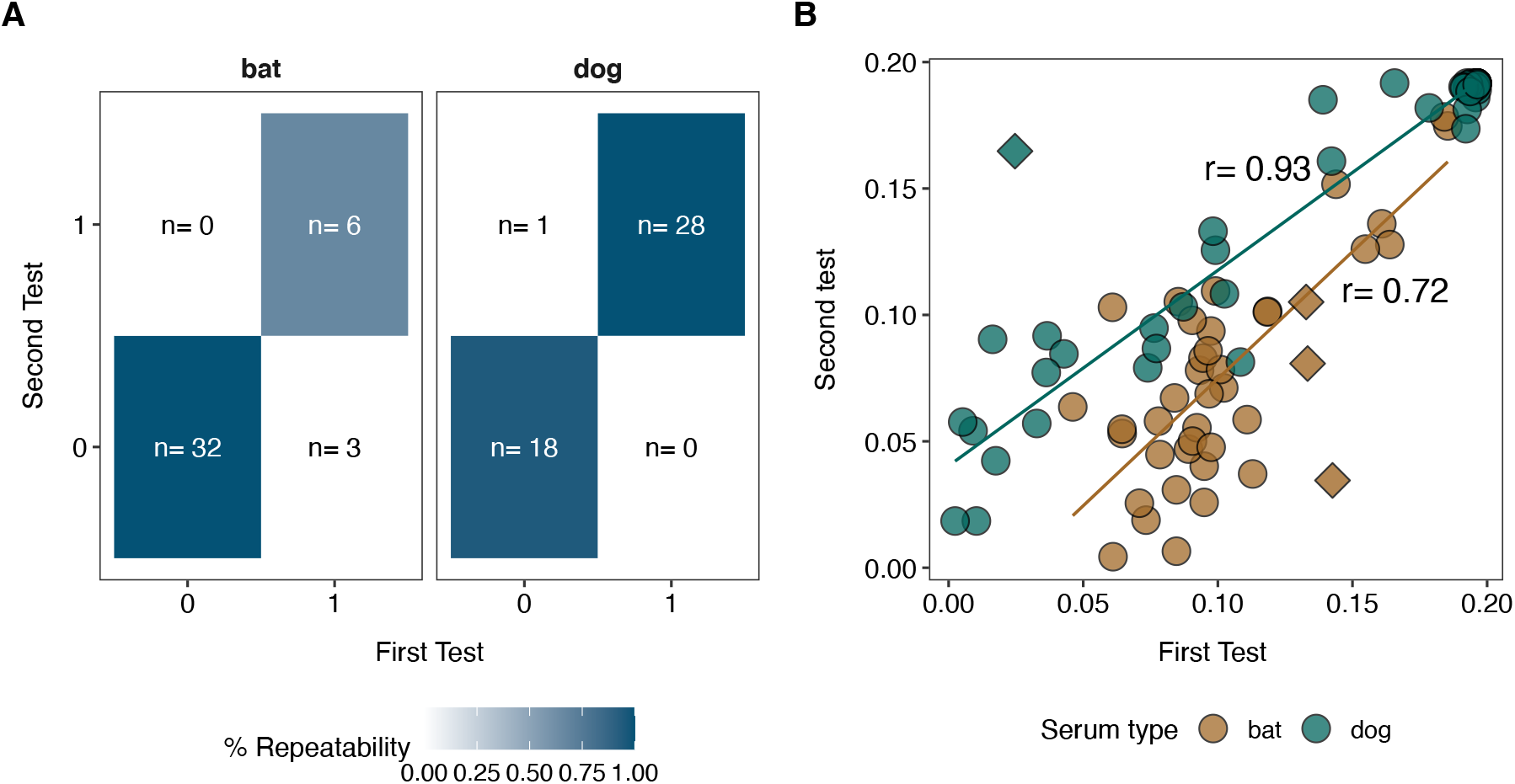
Repeatability of the predictions in bat (N = 41) and dog (N = 47) samples. A) Percentage of repeatability of the seropositive outcome of the bat and dog serum samples with the pmRFFIT test. B) Linear correlation of the predicted titers between the first and second test for dog serum and bat serum. Diamonds indicate samples that were contradictory in the binomial prediction.

Relative to FAVN, pmRFFIT predictions using the binomial model on dog sera showed overall moderate accuracy (80.43% of samples in agreement with FAVN; sensitivity = 78.79%; specificity = 84.62%). Accuracy was higher for the dog samples categorized as positive according to the binomial prediction (N = 28), where only 2 samples failed in the pmRFFIT prediction compared to the FAVN. For dog samples categorized as negative by the binomial pmRFFIT (N = 9), 7 samples were contradictory. These misclassified samples were consistently predicted as negative in the pmRFFIT, and titers displayed a strong correlation between tests (r = 0.93, SI 4). The binomial model was most accurate for samples with high RVNA titers (> 1.04 IU/mL, N = 13; 100% predicted to be seropositive; **Figure 5** A). At lower FAVN titers (< 1.04 IU/mL, N = 33), 72.73% of samples were predicted correctly by the pmRFFIT. To evaluate the accuracy of the binomial prediction in this lower range, the titer categorization process was repeated using the subset of data with FAVN titers ≤ 0.871 IU/mL (N = 33). Among the higher titer samples in this particular set (> 0.27 to ≤ 0.871 IU/mL, N = 11), 90.91% were accurately predicted as seropositive (**Figure 5** B). From the samples with titers ≤ 0.27 IU/mL, only 63.64% were accurately predicted (N = 22, **Figure 5** B). Accuracy rose to 83.33% in the lowest FAVN titer range (≤ 0.07 IU/mL, N = 12). The ROC curve showed that threshold values between 0.099 and 0.166 IU/mL maximized sensitivity and specificity.

**Figure 5.**
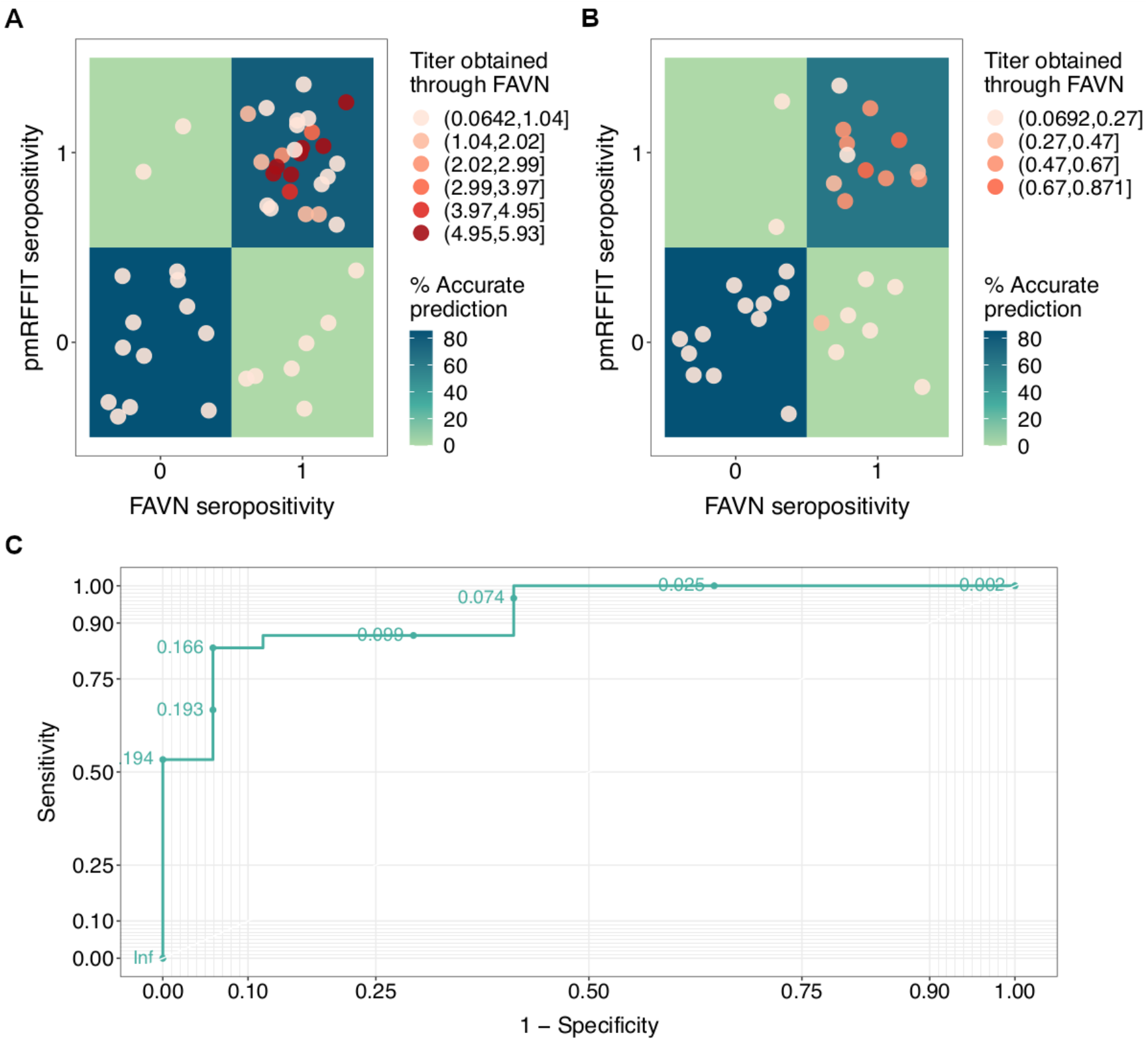
Validation of pmRFFIT with the FAVN on the dog serum samples. A) Seropositivity comparison for all FAVN titers: Sensitivity = 78.79%, Specificity = 85.71%, Overall accuracy = 80.85% (N = 47). B) Seropositive comparison of lower FAVN titers: Overall accuracy = 72.73% (N = 33). C) ROC curve of the lognormal titer prediction of the first test compared to the FAVN at threshold > 0.1 IU/mL for seropositivity. The continuous thresholds for the pmRFFIT are in green.

## Discussion

The most commonly applied serological tests to detect RVNA titers challenge a range of serial dilutions of serum with infectious RV. This process is labor-intensive and requires laboratory capacity to grow large quantities of pathogenic RV. Here, we provide an alternative serological framework that uses a combination of digital image analysis and statistical analysis to estimate the presence and titer of RVNA from a single dilution using only 3.5 μL of serum.

The pmRFFIT differs from other lyssavirus neutralization tests in several key aspects. It uses an MLV(RG) pseudotype rather than pathogenic RV, allowing the pmRFFIT to be performed in any low-containment laboratory with appropriate cell culture and microscopy facilities. The addition of GFP expression is significant, since it removes the need for FITC-conjugated antibody (reducing reagent costs) and the fixation and staining steps used in traditional RFFIT or FAVN. One potential drawback of using GFP expression to measure infectivity is the prolonged neutralization period (66 h versus 24 h RFFIT and 48 h FAVN) required to gain sufficient fluorescence for image processing (Aubert, 1992; Smith et al., 1973). Longer neutralization requirements (60 h) were also required in a FAVN modification using a GFP expressing recombinant CVS-11-eGFP but did not alter results relative to the test run with CVS-11 (Xue et al., 2014). Fortunately, extended incubations are unlikely to alter neutralization outcomes since MLV(RG) pseudotype is replication incompetent, preventing infection of additional cells during the incubation (Temperton et al., 2015). The pmRFFIT also uses an imaging pipeline that combines systematic photography of microscope fields with automated digital image processing to count infected cells. Microscopy in neutralization tests is time consuming and presents challenges for interlaboratory comparisons due to multiple sources of variation, especially those that affect the manual readout (e.g. laboratory user, manual pipetting, uneven cell monolayer) (Briggs et al., 1998; Cliquet et al., 1998; Eschbaumer et al., 2014; Péharpré et al., 1999; Timiryasova et al., 2019). The pmRFFIT standardized approach minimizes these sources of error while potentially reducing microscope operator time. Moreover, the imaging process generates traceable and permanent electronic records of the raw data, eliminating the need to manually digitize records of field counts. Several investigators have previously incorporated image processing into RV neutralization tests. For example, Pérhapré et al. (1999) modified the FAVN and RFFIT readout by using an automated image analysis, however one important drawback was the high cost of the equipment (a motorized stage and the software required for this). Streicker et al. (2012) also used photography and image processing to count pixels using a microRFFIT but did not make full use of the quantitative nature of imaging data to obtain RVNA titers and used pathogenic RV rather than a viral pseudotype. A final distinction is that instead of scoring microscope field or wells as virus positive or negative, the pmRFFIT predicts serological status and RVNA titer from infected cell counts in a single serum dilution using statistical modelling. The efficacy of this approach highlights the value of historically underutilized quantitative data on cellular infectivity for lyssavirus serology.

Model selection indicated substantial day-to-day variation in the SRIG dilution series. This was unsurprising, since virus neutralization tests are biologically dynamic systems that can be influenced by many factors (e.g. variability in the humidity of the incubator, technical manipulation, light condition of the microscope, variability in GFP expression in the cells) (Briggs et al., 1998; Hammami et al., 1999; Kostense et al., 2012). Since our statistical approach handles this variability through the random effect of test date, the pmRFFIT is best suited for large numbers of serum samples that require testing to be carried out across multiple batches. However, performance is only marginally reduced when running single models for each test date, implying the pmRFFIT may still be useful when fewer samples are available for testing (see SI 5, 6). Surprisingly, fitting the GLMMs to data from a single 1:10 dilution of SRIG predicted both seropositivity and RVNA titer more accurately than models fit to both the 1:10 and 1:25 dilutions. The reduced performance of the two-dilution model reflected higher variability in the 1:25 dilution compared to the 1:10 dilution, as evidenced by greater IQR values (SI 2). Ultimately, this variability likely reflects both higher stochasticity in infected cell counts at lower serum concentrations and pipetting error. Regardless, the ability to detect low titers (< 0.2 IU/mL) with just one dilution and without the need to conduct separate tests for screening and titration is advantageous since manual effort and materials are reduced.

Cell-based experiments are expected be more variable than other serological diagnostics, and often the allowed variation for test precision is up to 30% (Kostense et al., 2012; Timiryasova et al., 2019).

For example, Kostense et al. (2012) evaluated RFFIT repeatability by testing 3 different types of immunoglobulins (HRIG, CL184 and SRIG), and intra-test variation was 26%, 18% and 25% respectively. Similarly, Timiryasova et al. (2019) validated a RFFIT protocol with 15.7% variation among intra-test repetitions. In comparison, our test showed higher repeatability for serum samples from dogs and similar repeatability for serum samples from bats. Positive prediction for bat samples had a lower repetition, but most of the conflicts occurred with titers near the threshold defining seropositivity (2/3 samples, **Figure 4** B, SI 3) challenging a repeatable prediction between tests. The higher repeatability in dog sera relative to bat sera is likely related to the presence of noticeably higher neutralizing antibody titers in the dog sera, with 26/47 dogs (most of which had been vaccinated), compared to 5/41 bat samples having high predicted titers (≥ 0.15 IU/mL). Consistent with our findings, Kostenze et al. (2012) also observed that antibody concentration curves had an initial and evident bend in the titer concentration at 0.2 IU/mL, but the ability to observe slope changes in the lowest concentrations of serum (< 0.1 IU/ml) was challenging. Overall, the pmRFFIT was most precise in discriminating strong positives from strong negatives, where the highest and lowest titers were constantly predicted as such. In consideration of these points, standard serological methods might have limitations measuring low RVNA titers (< 0.5 IU/mL) while the pmRFFIT grants a highly repeatable approach for these titer values.

Using the external validation set of dog samples with known RVNA titers, the overall accuracy of the pmRFFIT was moderately high. Discrepancies occurred in samples with FAVN titers ranging from 0.07 to 0.29 IU/mL but were not observed for samples with higher titers (N = 23). In the previous analysis of these samples, sera that did not reach the 0.5 IU/mL threshold were considered negative and were not re-tested (Wright et al., 2009). We therefore cannot disregard the possibility of inaccurate titer assessment through FAVN, considering low-titer detection is even more challenging through classical approaches and that titer prediction through FAVN tests can vary even for higher titers (i.e. samples scoring 0.5 IU/mL can range from 0.38 to 0.66 IU/mL when repeated) (Liu, Zhang, Zhang, & Hu, 2012; Wright et al., 2009). Although we did not carry out a formal validation for the bat samples, the estimated seroprevalence was 0.21 (CI: 0.09, 0.35), consistent with several previous studies using the RFFIT (de Thoisy et al., 2016; Steece & Altenbach, 1989; Streicker, Franka, Jackson, & Rupprecht, 2013; Streicker et al., 2012). The ability of our approach to estimate VNA titers for all samples tested without additional laboratory work provides an additional layer of information that can be further used to gauge confidence in positive or negative predictions. For example, predicted titers abutting the threshold for seropositivity might be viewed with caution.

Detection of low RVNA titers is important for epidemiological studies of rabies exposure (Gold, Donnelly, Nouvellet, & Woodroffe, 2020). For example, bats sometimes produce low levels of RVNA after viral exposures, and the antibody response can wane to undetectable levels within months after the first exposure (Jackson et al., 2008; Turmelle et al., 2010). Indeed, the predicted titers among our predicted seropositive bat samples were consistently less than < 0.12 IU/mL. For this reason, we designed our test to detect low RVNA titers, defined as > 0.1 IU/mL, and the ROC analysis showed this was within the optimal range of thresholds for sensitivity and specificity. Consequently, titers > 0.2 IU/mL, as might be required for studies concerned with protective levels following immunization, cannot currently be estimated accurately. By pmRFFIT, such samples would be classified as seropositive with a predicted titer of 0.2 IU/mL. If desired, predicting higher RVNA titers than the focus of this study would require higher additional serum dilutions in conjunction with a greater range of SRIG concentrations.

In summary, the pmRFFIT quantifies RVNA titers using a single, low-biocontainment test that requires only a single dilution of test sera. We recommend first employing the binomial model at a predetermined threshold for seropositivity, then using the log-normal model to predict titers for the putatively positive samples (considered positive by the binomial model). If benchmark data are available, it would be possible to select the most sensitive and specific threshold after predicting titers for all available samples using the ROC curve analysis. Downstream analyses could use binomial and continuous data as needed for specific project objectives. The pmRFFIT could be extended to other lyssaviruses for similarly low-biocontainment testing of neutralizing antibodies. The ability to perform the test using the low-volume sera that are often available for studies of wild bats or longitudinal studies of captive bats is a particularly useful feature of our approach. Such studies are increasingly desirable to investigate the transmission dynamics of poorly understood bat viruses.

## Acknowledgments

This work was funded by a Sir Henry Dale Fellowship jointly funded by the Wellcome Trust and Royal Society (102507/Z/13/Z) and a Wellcome Senior Research Fellowship (217221/Z/19/Z). Additional funding was provided by the National Science Foundation (grant: DEB-1020966). DKM was funded by the Human Frontier Science Program (grant: RGP0013/2018) and the Mexican National Council for Science and Technology (CONACYT). We thank Carlos Tello for sampling coordination. We thank Carlos Shiva, Nestor Falcon, William Valderrama, Sergio Recuenco and John Claxton for assistance with sample storage and permits. We thank Pablo Murcia, Megan Griffiths, Nardus Mollentze and Laura Bergner for feedback on experimental design and earlier versions of this manuscript. SERFOR authorized the collection and exportation of samples (0142-2015 SERFOR/DGGSPFFS, N¤003327-SERFOR).

## Ethical statement

The authors confirm that the ethical policies of the journal, as noted on the journal’s author guidelines page, have been adhered to and the appropriate ethical review committee approval has been received. Bat capture and sampling methods were approved by the Research Ethics Committee of the University of Glasgow School of Medical Veterinary and Life Sciences (Ref 08a/15).

