## Supplemental material for "Predicting the presence and titer of rabies virus neutralizing antibodies from low-volume serum samples in low-containment facilities"

**Supplementary material**

**Sample information**

Supplementary material 1 Sample ID and information of collection

| ID 1 | Serum ID | Serum type | month | year | City | Country |
| --- | --- | --- | --- | --- | --- | --- |
| 3299 | SR1479 | bat | 5 | 2017 | Lima | Peru |
| 6613 | SR1426 | bat | 4 | 2017 | Lima | Peru |
| 6621 | SR1432 | bat | 4 | 2017 | Lima | Peru |
| 6638 | SR1468 | bat | 5 | 2017 | Lima | Peru |
| 7660 | SR1433 | bat | 4 | 2017 | Lima | Peru |
| 7662 | SR1430 | bat | 4 | 2017 | Lima | Peru |
| 7669 | SR1460 | bat | 4 | 2017 | Lima | Peru |
| 7671 | SR1436 | bat | 4 | 2017 | Lima | Peru |
| 7672 | SR1440 | bat | 4 | 2017 | Lima | Peru |
| 7686 | SR1456 | bat | 4 | 2017 | Lima | Peru |
| 8053 | SR1473 | bat | 5 | 2017 | Lima | Peru |
| 8527 | SR1392 | bat | 3 | 2017 | Lima | Peru |
| 8557 | SR1425 | bat | 4 | 2017 | Lima | Peru |
| 8568 | SR1462 | bat | 4 | 2017 | Lima | Peru |
| 8580 | SR1464 | bat | 4 | 2017 | Lima | Peru |
| 8588 | SR1448 | bat | 4 | 2017 | Lima | Peru |
| 8593 | SR1459 | bat | 4 | 2017 | Lima | Peru |
| 8606 | SR1458 | bat | 4 | 2017 | Lima | Peru |
| 8623 | SR1429 | bat | 4 | 2017 | Lima | Peru |
| 8649 | SR1480 | bat | 5 | 2017 | Lima | Peru |
| 8658 | SR1483 | bat | 5 | 2017 | Lima | Peru |
| 8668 | SR1481 | bat | 5 | 2017 | Lima | Peru |
| 8688 | SR1471 | bat | 5 | 2017 | Lima | Peru |
| 8701 | SR1484 | bat | 5 | 2017 | Lima | Peru |
| 8711 | SR1451 | bat | 4 | 2017 | Lima | Peru |
| 8713 | SR1428 | bat | 4 | 2017 | Lima | Peru |
| 8718 | SR1439 | bat | 4 | 2017 | Lima | Peru |
| 8721 | SR1446 | bat | 4 | 2017 | Lima | Peru |
| 8722 | SR1427 | bat | 4 | 2017 | Lima | Peru |
| 8724 | SR1463 | bat | 4 | 2017 | Lima | Peru |
| 8733 | SR1453 | bat | 4 | 2017 | Lima | Peru |
| 8736 | SR1467 | bat | 4 | 2017 | Lima | Peru |
| 8742 | SR1435 | bat | 4 | 2017 | Lima | Peru |
| 8744 | SR1438 | bat | 4 | 2017 | Lima | Peru |
| 8746 | SR1441 | bat | 4 | 2017 | Lima | Peru |
| 8747 | SR1445 | bat | 4 | 2017 | Lima | Peru |
| 8748 | SR1447 | bat | 4 | 2017 | Lima | Peru |
| 8750 | SR1454 | bat | 4 | 2017 | Lima | Peru |
| 8763 | SR1472 | bat | 5 | 2017 | Lima | Peru |
| 8764 | SR1475 | bat | 5 | 2017 | Lima | Peru |
| 8767 | SR1482 | bat | 5 | 2017 | Lima | Peru |
| Continues… | | | | | | |
| BS 02 08 | 350 | dog | 6 | 2008 | Bisarara | Tanzania |
| BS 04 08 | 352 | dog | 6 | 2008 | Bisarara | Tanzania |
| BS 06 08 | 154 | dog | unknown | 2008 | Bisarara | Tanzania |
| BS 101 08 | 395 | dog | 6 | 2008 | Bisarara | Tanzania |
| BS 12 08 | 360 | dog | 6 | 2008 | Bisarara | Tanzania |
| BS 13 08 | 161 | dog | unknown | 2008 | Bisarara | Tanzania |
| BS 13 08 | 361 | dog | 6 | 2008 | Bisarara | Tanzania |
| BS 147 08 | 397 | dog | 6 | 2008 | Bisarara | Tanzania |
| BS 16 08 | 164 | dog | unknown | 2008 | Bisarara | Tanzania |
| BS 20 08 | 166 | dog | 6 | 2008 | Bisarara | Tanzania |
| BS 22 08 | 368 | dog | 6 | 2008 | Bisarara | Tanzania |
| BS 68 08 | 185 | dog | unknown | 2008 | Bisarara | Tanzania |
| BS 75 08 | 183 | dog | 6 | 2008 | Bisarara | Tanzania |
| BS 76 08 | 386 | dog | 6 | 2008 | Bisarara | Tanzania |
| BS 77 08 | 184 | dog | unknown | 2008 | Bisarara | Tanzania |
| BS 77 08 | 384 | dog | 6 | 2008 | Bisarara | Tanzania |
| BS 80 08 | 381 | dog | 6 | 2008 | Bisarara | Tanzania |
| BS 82 08 | 180 | dog | unknown | 2008 | Bisarara | Tanzania |
| BS 83 08 | 178 | dog | 6 | 2008 | Bisarara | Tanzania |
| BS 87 08 | 188 | dog | unknown | 2008 | Bisarara | Tanzania |
| BS 91 08 | 391 | dog | 6 | 2008 | Bisarara | Tanzania |
| NG 117 08 | 29 | dog | unknown | 2008 | Unknown | Tanzania |
| NG 123 08 | 230 | dog | unknown | 2008 | Unknown | Tanzania |
| NG 144 08 | 236 | dog | unknown | 2008 | Unknown | Tanzania |
| NG 225 08 | 45 | dog | unknown | 2008 | Unknown | Tanzania |
| NG 24 08 | 212 | dog | unknown | 2008 | Unknown | Tanzania |
| NY 04 08 | 117 | dog | 5 | 2008 | Nyamburi | Tanzania |
| NY 05 08 | 102 | dog | 5 | 2008 | Nyamburi | Tanzania |
| NY 104 08 | 317 | dog | 5 | 2008 | Nyamburi | Tanzania |
| NY 113 08 | 23 | dog | 5 | 2008 | Nyamburi | Tanzania |
| NY 117 08 | 126 | dog | 5 | 2008 | Nyamburi | Tanzania |
| NY 177 08 | 132 | dog | 5 | 2008 | Nyamburi | Tanzania |
| NY 179 08 | 134 | dog | 5 | 2008 | Nyamburi | Tanzania |
| NY 183 08 | 140 | dog | 5 | 2008 | Nyamburi | Tanzania |
| NY 189 08 | 146 | dog | 5 | 2008 | Nyamburi | Tanzania |
| NY 192 08 | 136 | dog | 5 | 2008 | Nyamburi | Tanzania |
| NY 195 08 | 139 | dog | 5 | 2008 | Nyamburi | Tanzania |
| NY 20 08 | 106 | dog | 5 | 2008 | Nyamburi | Tanzania |
| NY 200 08 | 137 | dog | 5 | 2008 | Nyamburi | Tanzania |
| NY 22 08 | 108 | dog | 5 | 2008 | Nyamburi | Tanzania |
| NY 54 08 | 109 | dog | 5 | 2008 | Nyamburi | Tanzania |
| NY 60 08 | 114 | dog | 5 | 2008 | Nyamburi | Tanzania |
| NY 67 08 | 116 | dog | 5 | 2008 | Nyamburi | Tanzania |
| NY 91 07 | 148 | dog | 5 | 2008 | Nyamburi | Tanzania |
| RNG 03 08 | 53 | dog | 5 | 2008 | Rungabure | Tanzania |
| Continues… | | | | | | |
| RNG 26 08 | 68 | dog | 5 | 2008 | Rungabure | Tanzania |
| RNG 264 08 | 97 | dog | 5 | 2008 | Rungabure | Tanzania |

**Variability in the counts**

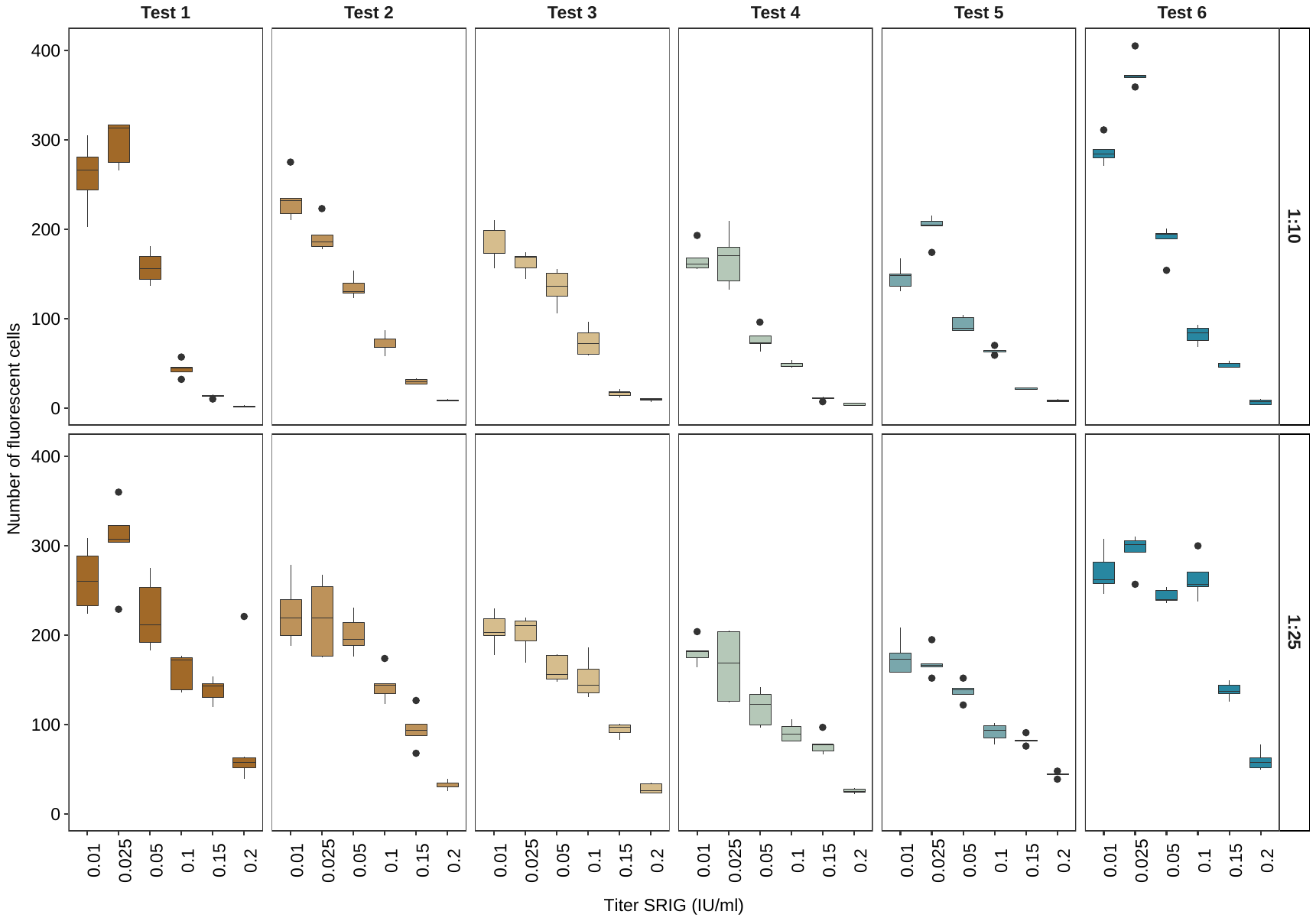

Supplementary material 2. Variability of the interquartile ranges of the SRIG concentrations

**Repeatability between tests, inconsistent samples**

Supplementary material 3 Inconsistent samples between the first and second pmRFFIT binomial predictions with a threshold >0.1 IU/mL and their respective pmRFFIT titer predictions

| Serum ID | Serum type | 1^st^ binomial prediction | 2^nd^ binomial prediction | 1^st^  titer prediction  (IU/mL) | 2^nd^  titer prediction  (IU/mL) |
| --- | --- | --- | --- | --- | --- |
| SR1484 | bat | 1 | 0 | 0.143 | 0.035 |
| SR1451 | bat | 1 | 0 | 0.133 | 0.105 |
| SR1446 | bat | 1 | 0 | 0.133 | 0.081 |
| 114 | dog | 0 | 1 | 0.025 | 0.165 |

*Validation pmRFFIT vs FAVN, contradictory results*

Supplementary material 4 Inconsistent samples between the first pmRFFIT binomial predictions and the original binomial result from FAVN (both considering as threshold >0.1 IU/mL), with their respective titer values obtained through pmRFFIT and FAVN

| Serum ID | Serum type | 1^st^ pmRFFIT  binomial prediction | FAVN  binomial result | 1^st^ pmRFFIT  titer prediction  (IU/mL) | 2^nd^ pmRFFIT  titer prediction  (IU/mL) | Titer obtained through FAVN  (IU/mL) |
| --- | --- | --- | --- | --- | --- | --- |
| 164 | dog | 0 | 1 | 0.04 | 0.08 | 0.29 |
| 178 | dog | 0 | 1 | 0.09 | 0.10 | 0.13 |
| 29 | dog | 0 | 1 | 0.07 | 0.08 | 0.17 |
| 45 | dog | 0 | 1 | 0.03 | 0.06 | 0.13 |
| 23 | dog | 0 | 1 | 0.10 | 0.13 | 0.13 |
| 68 | dog | 0 | 1 | 0.08 | 0.09 | 0.17 |
| 146 | dog | 0 | 1 | 0.08 | 0.09 | 0.17 |
| 137 | dog | 1 | 0 | 0.19 | 0.19 | 0.07 |
| 108 | dog | 1 | 0 | 0.14 | 0.16 | 0.07 |

**Analysis with GLMs for each individual test date**

To evaluate results on single test dates a generalized linear model (GLM) was perform for each individual test. Every single GLM was independently fitted to the single SRIG dilution curve from the individual date. The models built with only the 1:10 dilution data performed much better than models build with both the 1:10 and 1:25 dilution data. Similar to predictions from the GLMM analyses, both dilution models had equivalent specificity (100%), but the one-dilution model was more sensitive (100% versus 58.33%). Observed and predicted SRIG titers were higher correlated for the one-dilution model overall and in individual test dates. When comparing the predictions of the GLM versus the GLMM across tests, 6 out of 245 predictions were contradictory (3.8%, 4.88%, 0%, 4.88%, 0%, 2.13% respectively for each direct comparison). The correlation of the titer prediction was high for all the comparisons (r>=0.99). Note that the GLM predicted some negative titer values, this might be related to the numbers of fluorescent cells in the sample being much higher than the lowest control concentration in the analysis (0.05 IU/mL). For this scenario the sample should be assumed to have a titer concentration <0.05 IU/mL.

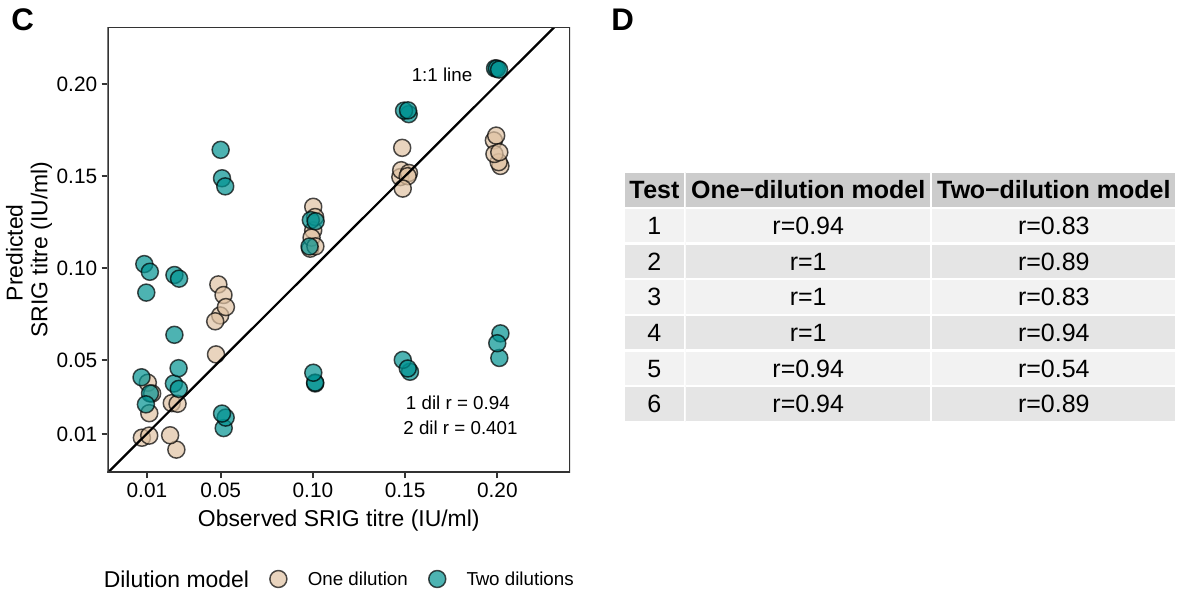

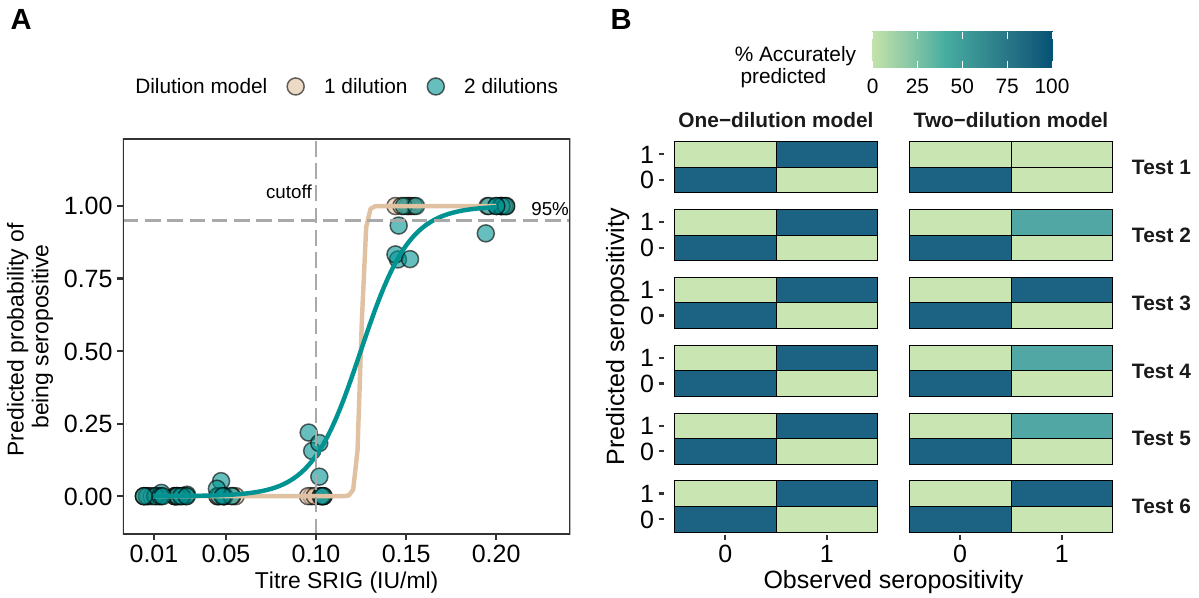

Supplementary material 5 Results of the independent GLM models fit to the SRIG control data. All graphs compare the one-dilution (1:10) and the two-dilution (1:10 and 1:25) models.

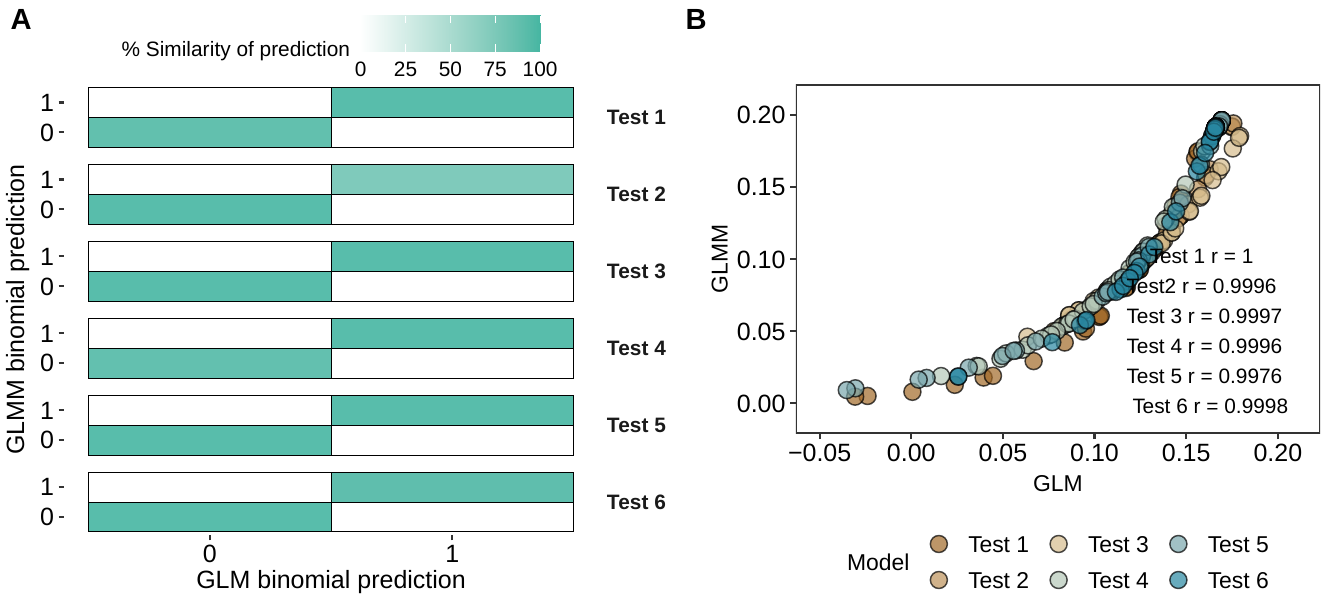

Supplementary material 6 Comparison of the results between the GLM and the GLMM predictions across all the tests
